# Cannabidiol as a Prophylactic Agent Against Glioblastoma Growth: A Preclinical Investigation

**DOI:** 10.1101/2025.10.29.685440

**Authors:** Lei P Wang, Bidhan Bhandari, Sahar Emami Naeini, Breanna Hill, Hannah M Rogers, Jules Gouron, Nayeli Perez-Morales, Aruba Khan, William Meeks, Ahmed El-Marakby, Nancy Young, Fernando L Vale, Salman Ali, Gerald Wallace, Jack C Yu, Ali S Arbab, Évila Lopes Salles, Babak Baban

## Abstract

**Purpose:** Glioblastoma (GBM) is one of the most aggressive brain tumors, with limited treatment options and poor outcomes due to frequent relapse after surgery. This study aims to investigate whether pretreatment with inhaled cannabidiol (CBD) can inhibit GBM growth in a preclinical murine model. Specifically, we hypothesize that CBD pretreatment may reduce tumor progression and modulate the tumor microenvironment.

**Methods:** C57BL/6 mice were pretreated with inhaled CBD for either 3 or 14 days, or a placebo, followed by intracranial implantation of glioblastoma cells. Tumor growth, immune checkpoint expression (IDO, PD-L1), and key biomarkers (MGMT, Ki67) were assessed to determine the impact of CBD pretreatment on tumor progression and the immune microenvironment.

**Results:** The 14-day CBD pretreatment significantly reduced tumor growth compared to both the placebo and 3-day CBD groups. Additionally, this group showed lower expression of immune checkpoints (IDO, PD-L1) and reduced levels of MGMT and Ki67, suggesting enhanced tumor suppression. These results indicate that prolonged CBD pretreatment modulates the tumor microenvironment and may improve tumor control and reduce relapse risk.

**Conclusion:** This study demonstrates that inhaled CBD pretreatment significantly inhibits GBM growth in a preclinical model. CBD’s ability to reduce immune checkpoint expression and key biomarkers associated with tumor progression suggests that it could be an effective strategy for enhancing the efficacy of GBM therapies and potentially improving post-surgical outcomes. Further research is required to investigate its clinical potential and mechanisms of action.

## Introduction

Glioblastoma (GBM) is the most prevalent and aggressive primary malignant brain tumor in adults, characterized by rapid proliferation, extensive infiltration, and a dismal prognosis. Despite decades of research and therapeutic advances, the median survival for GBM patients remains approximately 15 months following diagnosis, with minimal improvement over time [1–4]. Since 1926, only a limited number of pharmacological agents and a single medical device have received regulatory approval for GBM treatment, underscoring the persistent therapeutic challenges associated with this disease [5]. The etiology of GBM remains poorly understood, and its rising incidence further emphasizes the urgent need for innovative therapeutic and preventive strategies [6].

Emerging research suggests that early intervention or prophylactic approaches may significantly alter the trajectory of GBM progression. Preventative strategies, particularly those targeting molecular pathways involved in tumor initiation and resistance, hold promise for reducing tumor burden, delaying neurological deterioration, and improving overall survival [7–9]. Shifting the treatment paradigm from reactive to proactive necessitates a focus on agents capable of modulating GBM-associated pathophysiology before overt tumor development.

Cannabidiol (CBD), a non-psychoactive phytocannabinoid derived from Cannabis sativa, has attracted growing interest for its broad spectrum of therapeutic properties, including anti-cancer and neuroprotective effects [1,3]. Preclinical studies have demonstrated that CBD exerts anti-proliferative, pro-apoptotic, anti-inflammatory, and anti-angiogenic effects across various tumor models, including GBM [3,10–12]. Our previous work and other studies have shown that CBD can inhibit GBM growth in the brain by modulating the endocannabinoid system, promoting cell cycle arrest, and impairing angiogenesis [3,10–12]. These mechanisms suggest that CBD may be uniquely suited to address both the malignant and neurological aspects of GBM. Moreover, CBD’s established safety profile, lack of psychoactive effects, and reported neuroprotective and anti-inflammatory properties further support its suitability as a preventive agent in at-risk populations.

Notably, the prophylactic potential of inhaled CBD in GBM remains unexplored. Inhalation offers several pharmacokinetic advantages, including rapid systemic absorption, non-invasiveness, and efficient central nervous system delivery via enhanced blood-brain barrier (BBB) penetration [3,10,13]. Furthermore, CBD has been shown to downregulate O6-methylguanine-DNA methyltransferase (MGMT), a DNA repair enzyme associated with resistance to temozolomide, the current standard-of-care chemotherapy for GBM, further highlighting its potential as a sensitizing agent [14–15].

In this study, we investigate the prophylactic potential of chronic CBD administration via inhalation in a murine model of GBM, a novel approach that, to our knowledge, has not been previously explored. While CBD has been studied primarily in the context of treatment following tumor establishment, its utility as a preventive agent represents a significant departure from conventional paradigms focused solely on tumor eradication. The use of inhalation as a delivery method offers distinct translational advantages, including non-invasiveness, rapid systemic uptake, and efficient central nervous system penetration, making it particularly well-suited for potential clinical application in at-risk or pre-symptomatic populations. We hypothesize that sustained pretreatment with inhaled CBD will attenuate tumor growth by inducing apoptosis, suppressing cell proliferation, and downregulating chemoresistance mechanisms such as MGMT expression.

Demonstrating the efficacy of this approach could not only establish a foundation for preventive neuro-oncology strategies but also support the clinical development of CBD as a safe, non-toxic, and accessible adjunct to current GBM interventions. If validated, this work would position inhaled CBD as a paradigm-shifting strategy with the potential to delay disease onset, improve treatment responsiveness, and ultimately extend survival in patients at risk for or in the early stages of GBM.

## Materials and methods

### Animals

Wild-type (total of 30 mice from 2 independent cohorts, n=5 for each experimental group), 12 week old male mice (obtained from Jackson Laboratories, Bar Harbor, ME) were used to generate the orthotopic GBM model. The animals were housed in the laboratory animal facilities of the Augusta University with free access to food and water. All experiments were conducted under the approval of the Augusta University Animal Care and Use Committee (Protocol # 2011-0062).

### Metered aerosolized cannabinoid delivery device

The metered dose tincture inhaler used in this study, ApelinDx, was generously supplied by Thriftmaster Global Bioscience USA, as previously detailed [3]. For adaptation to the murine model, the device was customized by incorporating an additional nozzle component, allowing for better regulation of CBD delivery and inhalation volume. Each unit of ApelinDx contained a total of 1000 mg of formulation, comprising 985 mg of broad-spectrum cannabidiol (CBD) derived from winterized crude hemp extract, along with 15 mg of co-solvent, surfactant, and propellant. The device delivered 5 mg of CBD per spray at a controlled flow rate of 200 mL/min. The placebo formulation was identical in composition except that the active CBD component was substituted with 985 mg of hemp seed oil. Pharmacokinetic data indicate that inhaled CBD reaches peak plasma concentrations rapidly—typically within 3 to 10 minutes—achieving significantly higher maximum levels compared to oral delivery. Studies report an average systemic bioavailability of approximately 31% for inhaled CBD. Additionally, inhalation has been shown to lower levels of inactive circulating metabolites and enhance overall CBD absorption. These characteristics contribute to a more favorable therapeutic profile by ensuring consistent bioavailability, avoiding first-pass hepatic metabolism, and minimizing both systemic and localized adverse effects [3,16–17]. Regarding inhalation variability, the delivery was standardized through the use of a calibrated actuator and consistent environmental conditions within the exposure chamber. All animals received the inhaled dose from freshly primed canisters, ensuring uniformity across exposures. While minor variability is inherent to aerosolized delivery systems, our procedures were designed to minimize fluctuations in exposure across sessions.

### Pre-treatment with CBD inhalation

Mice were randomly allocated using simple randomization into three experimental groups (n=5/group): one control group and two CBD-treated groups. All mice were selected within a narrow weight range (28-29.5g) and weighed prior to treatment initiation to ensure consistent body-weight-adjusted dosing (based on average weight of 29 g). The CBD groups received six actuations of CBD (approximately 10 mg per animal) daily, starting on day two weeks (day −14) and three days (day −3) prior to tumor implantation in the brain. To minimize handling stress, mice were allowed to acclimate before CBD inhalation, with appropriate time intervals between the six actuations. These two distinct pre-treatment timeframes were chosen to assess the time-dependent effects of CBD on tumor progression. The dose, as used in our previous work, was calculated based on effectiveness and tolerability of CBD to achieve antitumor effect [3]. The control group was administered a placebo using a calibrated inhaler. As described previously [3], the inhalation procedure was conducted in a controlled environment to ensure accurate dosing and minimize variability. This pre-treatment protocol was designed to investigate the potential impact of CBD on modulating the tumor microenvironment and inhibiting tumor growth in the brain.

### Tumor cell Preparation and Orthotopic Glioblastoma Model in Mice

To establish the orthotopic glioblastoma (GBM) model, we followed previously validated protocols [3]. Luciferase-expressing GL261 murine glioma cells, which are syngeneic to C57BL/6 mice, were cultured in RPMI-1640 medium supplemented with 10% fetal bovine serum under standard conditions. On the day of implantation, cells were harvested and suspended in serum-free medium.

Mice were anesthetized with 3% isoflurane for induction and maintained at 1.5–2% throughout the surgical procedure. Following sterile preparation, a cranial burr hole was carefully drilled 2.25 mm lateral and 1 mm posterior to the bregma, ensuring the dura remained intact. A total of 30,000 GL261 cells suspended in 3 μL of media were loaded into a 10 μL Hamilton syringe fitted with a 26-gauge needle. The needle was inserted to a depth of 4 mm and then retracted to 3 mm, where the injection was performed. To ensure consistent and accurate delivery, all injections were performed using a motorized injector system, which allows for highly precise and reproducible administration of cells at the targeted depth and location. This standardized approach minimizes variability across experimental groups. To minimize cell reflux, the needle was withdrawn incrementally in 1 mm steps, beginning 2–3 minutes after cell delivery. The injection site was sealed with bone wax, and the exposed skull was disinfected using Betadine prior to skin closure with sutures. Postoperative pain management was provided via a single subcutaneous dose of buprenorphine (1 mg/kg). Tumor development was monitored on day 8 post-implantation using in vivo bioluminescence imaging. Mice received an intraperitoneal injection of D-luciferin (100 μL at 150 mg/kg), and bioluminescent signals were captured using the AmiX optical imaging system (Spectral Instruments Imaging, Tucson, AZ). Photon emission (photons/sec/mm^2^) was quantified using Aura imaging software (version 4.0.0; Spectral Instruments Imaging, LLC), enabling visualization of primary tumor burden and potential metastatic spread.

### Bioluminescence Imaging for Tumor Monitoring

To assess tumor burden and progression, bioluminescence imaging was performed on day 8 following intracranial tumor cell implantation. Mice received an intraperitoneal injection of D-luciferin (100 µL at a dose of 150 mg/kg), after which in vivo optical images were captured to evaluate both primary tumor growth and potential metastatic dissemination. Imaging was conducted using the AmiX optical imaging platform (Spectral Instruments Imaging, Tucson, AZ), and signal intensity was measured in photons per square millimeter per second. Quantitative analysis of the emitted light was carried out using Aura Imaging Software (version 4.0.0; Spectral Instruments Imaging, LLC).

To longitudinally monitor tumor development, additional imaging sessions were conducted on days 7 and 21 post-implantation. At the end of the imaging schedule, all animals were humanely euthanized, and brain tissues were harvested for downstream applications, including histopathological evaluation, immunofluorescence staining, and flow cytometric analyses.

### Visualization of Glioblastoma Lesions

To enable macroscopic examination of intracranial tumor growth, a craniotomy was performed on mice to expose the tumor-bearing region of the brain. Upon surgical exposure, high-resolution digital images were acquired to capture the gross morphology and visual characteristics of the tumor. This imaging approach facilitated direct comparison of tumor presentation between the CBD-pretreated and placebo-treated groups, offering real-time visual confirmation of differential tumor progression within the brain tissue.

### Histologic Analysis of Tumor Tissue

Histopathological analysis was performed following previously established protocols [1,3]. Glioblastoma tissues were freshly excised and fixed in 10% neutral buffered formalin (HT50-1-128; Sigma-Aldrich). Samples were then processed through standard dehydration steps and embedded in paraffin. All procedures were carried out at ambient room temperature. Paraffin-embedded brain sections were cut into 4 μm slices and stained with hematoxylin and eosin (H&E) using conventional histological techniques. Tissue morphology and tumor architecture were examined under a Zeiss brightfield microscope. Areas of tumor infiltration were measured and quantified using ImageJ software (version 1.53g; National Institutes of Health, Bethesda, MD, USA).

### Fluorescence-Based Immunohistochemical Analysis

Immunofluorescence staining was carried out on paraffin-embedded brain tumor sections using previously established protocols [3]. Tissue sections were incubated with fluorescently labeled primary antibodies targeting MGMT (O6-methylguanine-DNA methyltransferase; Novus Biologicals, USA, Cat# NB100-168SS) and the proliferation marker Ki-67 (Thermo Fisher Scientific, Cat# 12-5698-82). Nuclear counterstaining was performed with DAPI (4′,6-diamidino-2-phenylindole) to facilitate cellular visualization. Fluorescent images were acquired using a Zeiss fluorescence microscope. Quantitative image analysis was performed by selecting regions of interest (ROIs) using the lasso tool in Adobe Photoshop CS4 Extended. Within these ROIs, integrated density (pixel intensity) and mean gray value (a measure of fluorescence brightness) were recorded to assess relative marker expression levels.

### Flow Cytometric Analysis

For flow cytometry, a single-cell suspension was prepared from GBM tumor tissue by passing the sample through a 100 μm cell strainer (BD Biosciences, San Diego, CA, USA), followed by centrifugation at 252 g for 10 minutes. The resulting cell pellet was then subjected to a standard flow cytometry staining protocol as previously outlined [3]. Cells were fixed, permeabilized, and subsequently stained for the detection of intracellular signaling markers. Specific antibodies used included anti-SOX2 (APC-conjugated anti-mouse SOX2, R&D Systems, Cat# IC2018A), anti-PD-L1 (Alexa Fluor 488-conjugated anti-mouse PD-L1, Thermo Fisher Scientific, Cat# 53-5982-82), and anti-IDO (PerCP-conjugated anti-mouse IDO, Thermo Fisher Scientific, Cat# 46-9473-82). The stained cells were analyzed using a NovoCyte Quanteon flow cytometer (Agilent Technologies, Santa Clara, CA, USA), with data analysis performed using FlowJo V10 software. To validate antibody specificity and exclude potential non-specific binding to Fc receptors or other cellular components, appropriate isotype controls were included in all experiments. These isotype controls matched the primary antibody in terms of host species, isotype, and conjugation type, ensuring accurate results.

### Western blotting

Western blotting was used to assess the expression levels of MGMT in tumor tissues [18]. The samples were homogenized in RIPA lysis buffer in presence of protease inhibitor (Thermo Scientific). Protein concentration was determined using the BCA assay. Homogenates (50 μg protein) were separated by electrophoresis on a 4–15% precast polyacrylamide gel (Bio-Rad) and transferred to PVDF membrane that was previously soaked in methanol using PowerPac™ Universal power supply and Trans-Blot Turbo transferring system (Bio-Rad Laboratories, Inc.), respectively. Membranes were probed simultaneously with anti-mouse MGMT monoclonal antibody (O6-methylguanine-DNA methyltransferase, Novus Biologicals USA, Cat# NB100-168SS) and β-actin antibody (Thermo Fisher) for 24 hours. Proteins were detected using appropriate secondary antibodies conjugated with different fluorescent dyes compatible with the Odyssey imaging system, allowing simultaneous detection without stripping. Membranes were stripped and re-probed for β-actin (Thermo Fisher) to demonstrate equal loading. The reactive bands were visualized using the Li-Cor Odyssey FC system. The results were quantified by densitometry analysis utilizing ImageJ NIH software and expression levels were reported relative to β-actin.

### Statistical Analysis

To assess statistical significance between groups, we applied the Brown-Forsythe and Welch analysis of variance (ANOVA) with a significance threshold set at p < 0.05. For the quantification of tissue expression, comparisons between the placebo and CBD-treated groups were made using two-way ANOVA, followed by the post-hoc Sidak test to account for multiple comparisons (p < 0.05). Survival analysis was performed using the Kaplan-Meier method, with statistical differences between groups evaluated through the Log-rank Mantel-Cox test.

## Results

### Inhaled CBD Pretreatment Inhibits GBM Growth and Improves Tumor-Associated Health Indicators

In vivo bioluminescent imaging (BLI) performed at 7 and 21 days post-tumor implantation revealed that 14 days of inhaled CBD pretreatment markedly inhibited GBM tumor growth compared to placebo and 3-day CBD pretreatment groups (Figure 1A). Total photon emission quantification showed significantly reduced signal in the 14-day CBD group at both time points (*p < 0.05, ***p < 0.001), confirming decreased tumor burden (Figure 1B). Cross-validation of BLI with anatomical tumor volume (Figure 1C) further confirmed the reduced tumor burden, even in cases with modest increases in photon flux. To support translational relevance, body weight measurements indicated minimal weight loss in the 14-day CBD group (Figure 1D), suggesting improved systemic tolerance despite tumor progression, although not statistically significant. Survival analysis (Figure 1E), conducted across two independent cohorts (n = 10/group), showed significantly improved survival in the 14-day CBD group versus placebo (*p < 0.05). Notably, the BLI and imaging-based outcomes were derived from a representative cohort (n = 5/group) with full survival in CBD-treated animals, facilitating consistent longitudinal imaging.

**Figure 1.**
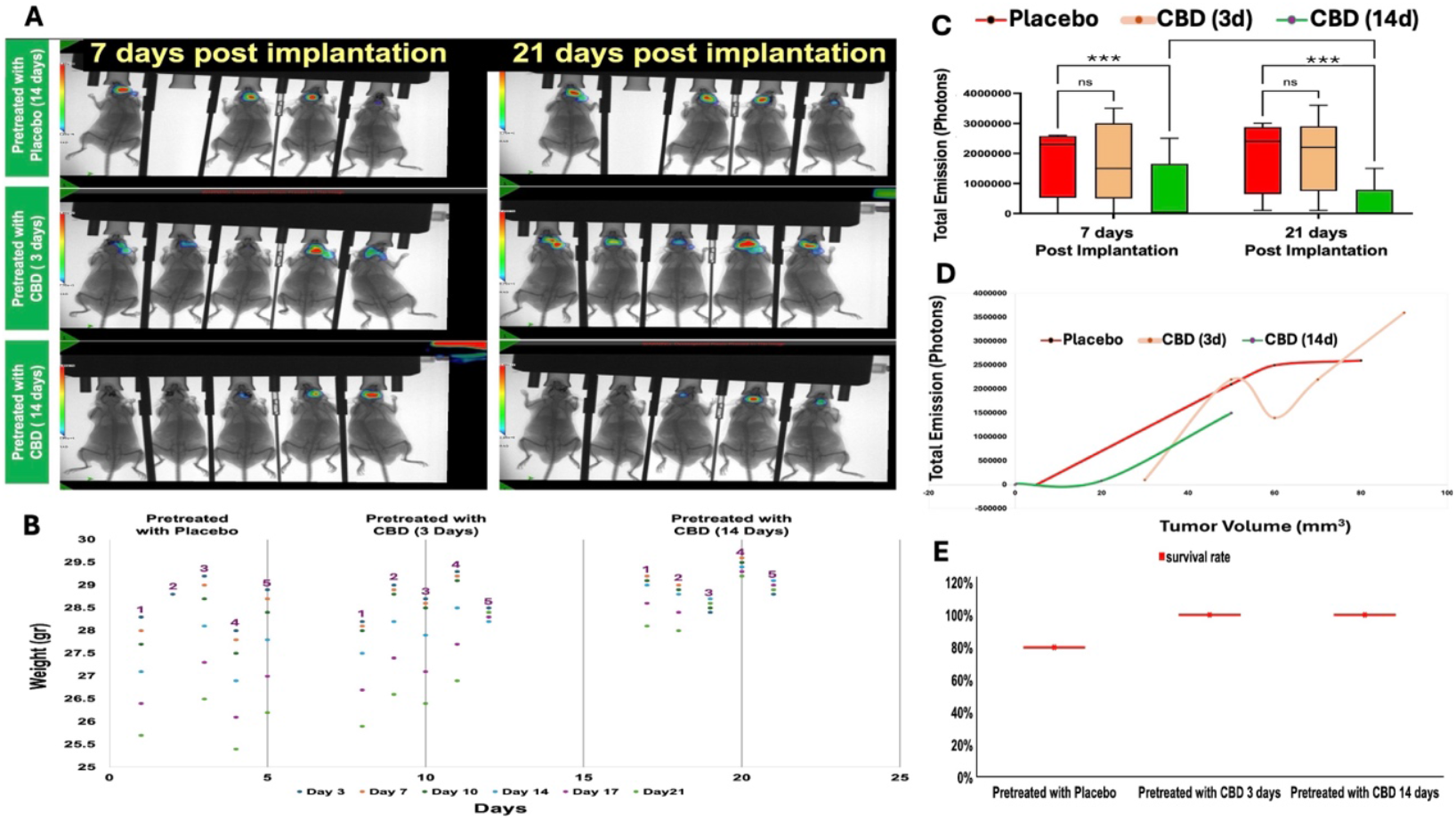
Inhaled CBD pretreatment inhibits GBM growth and improves associated health indicators. **A)** Optical bioluminescent imaging (BLI) of photon intensities at 7 and 21 days post–tumor implantation shows that 14 days of inhaled CBD pretreatment significantly reduced GBM tumor growth compared to both placebo and 3-day CBD pretreatment groups (p < 0.01). Tumors are indicated by yellow plus signs at the start of the study, red plus signs represent tumor growth, and green minus signs indicate tumor suppression or reduction. **B)** Quantification of total photon emission confirms significant tumor growth inhibition in the 14-day CBD group at both 7 and 21 days post-implantation (p < 0.05 and ***p < 0.001, respectively), while the 3-day pretreatment group did not differ significantly from placebo. **C)** Cross-validation of BLI signal with tumor volume measurements demonstrates consistent correlation between reduced photon flux and decreased tumor burden in the 14-day CBD group. This also confirms that despite the relatively flat BLI increase, true tumor growth differences were present. **D)** Body weight changes over time in all groups show minimal reduction in the 14-day CBD group, suggesting better systemic health, although differences were not statistically significant. **E)** Survival analysis across two cohorts (n=10/group) shows significantly increased survival in the 14-day CBD pretreatment group compared to placebo (p < 0.05). Survival analysis was performed across two cohorts (n=10/group) to enhance statistical power, while BLI and other outcomes were shown from one representative cohort (n=5/group) with full survival in CBD-treated groups for consistent imaging.

### CBD Pretreatment Reduces Tumor Volume and Improves Histopathology

Consistent with the imaging results, gross anatomical inspection and brain histology (Figure 2A) demonstrated visibly reduced tumor size in the 14-day CBD group compared to placebo and 3-day groups. Tumor volume quantification confirmed significantly smaller tumors in the 14-day CBD group (*p < 0.05; Figure 2B). Tumor area analysis also showed a marked reduction (***p < 0.001; Figure 2C). Histopathological evaluation revealed improved tumor architecture, reduced mitotic figures (yellow arrows), and lower levels of apoptosis (red arrows) in the 14-day group, suggesting that prolonged inhaled CBD pretreatment attenuates both tumor proliferation and cellular degeneration.

**Figure 2.**
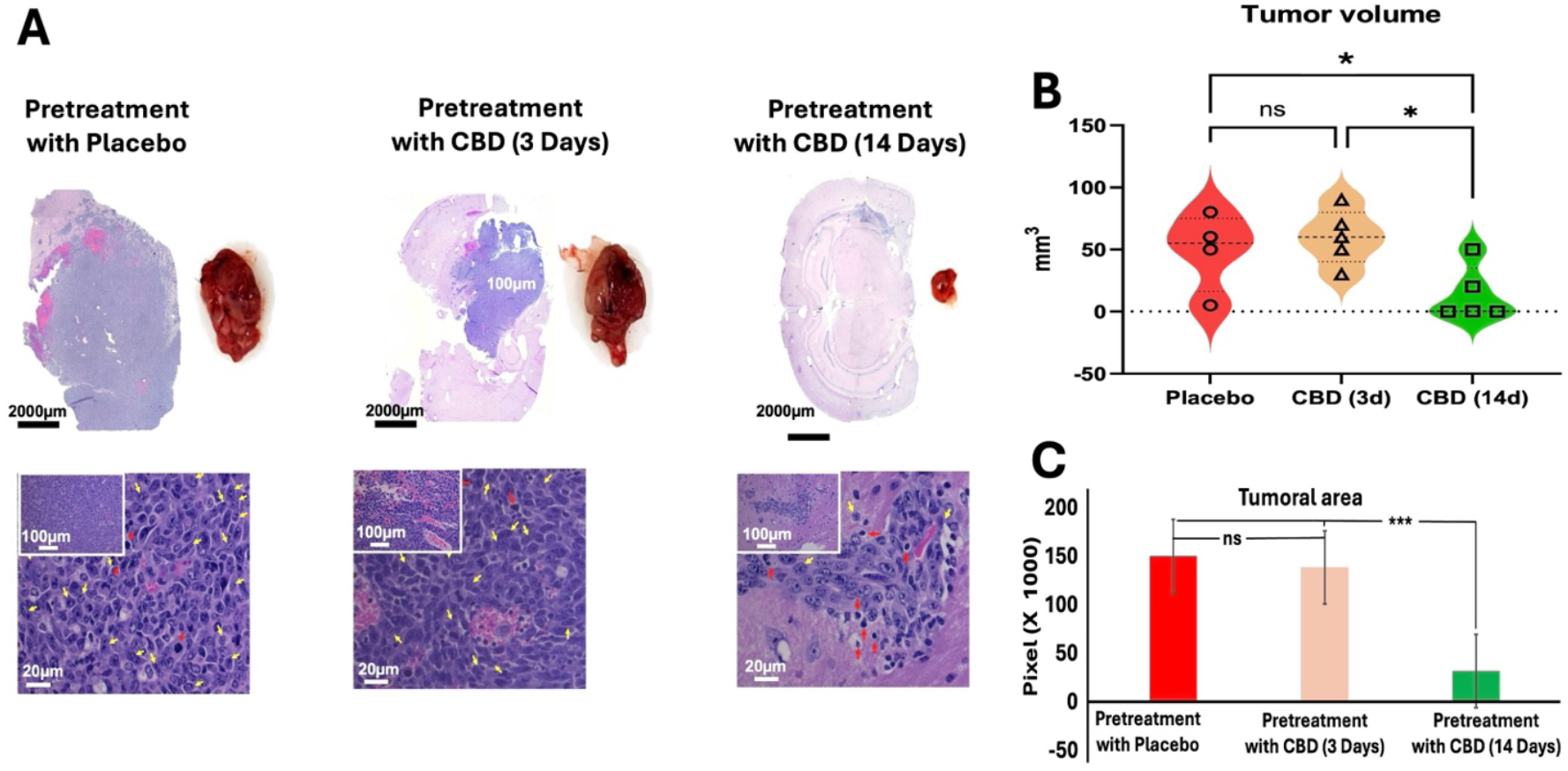
Prolonged Inhaled CBD Pretreatment Reduces Tumor Burden and Improves Tumor Histology. **A)** Representative brain sections and gross tumors from mice pretreated with placebo, 3-day CBD, or 14-day CBD show visibly reduced tumor size in the 14-day group. Scale bars: 2000 μm for whole brain sections; 100 μm and 20 μm for inset and main histology images, respectively. **B)** Quantified tumor volumes indicate a significant reduction in the 14-day CBD group compared to placebo and 3-day groups (p < 0.05). **C)** Tumor area analysis confirms these findings (p < 0.001). Histological examination reveals better tissue organization, lower mitotic activity (yellow arrows), and reduced apoptosis (red arrows) in the 14-day group.

### Inhaled CBD pretreatment reduces immune evasion gatekeepers in GBM

Mice pretreated with inhaled CBD for 14 days showed a significant reduction in the expression of immune evasion markers, IDO and PD-L1, in GBM tumors compared to both the placebo and 3-day CBD groups. Flow cytometry analysis (Figure 3A) identified tumor cells based on forward light scatter (FSC) and side light scatter (SSC) (left panels), followed by gating for SOX2 expression, a marker of stem-like properties (middle panels). Importantly, 14-day CBD pretreatment not only decreased overall SOX2 expression in the tumors but also specifically reduced IDO and PD-L1 expression within SOX2-positive cells (dot plots). Quantification of these markers in SOX2-positive cells (Figure 3B) revealed a significant decrease in both IDO and PD-L1 expression following 14-day CBD pretreatment (p < 0.001). These results suggest that prolonged CBD pretreatment may disrupt immune evasion mechanisms in GBM, particularly in stem-like cells, potentially enhancing anti-tumor immune responses.

**Figure 3.**
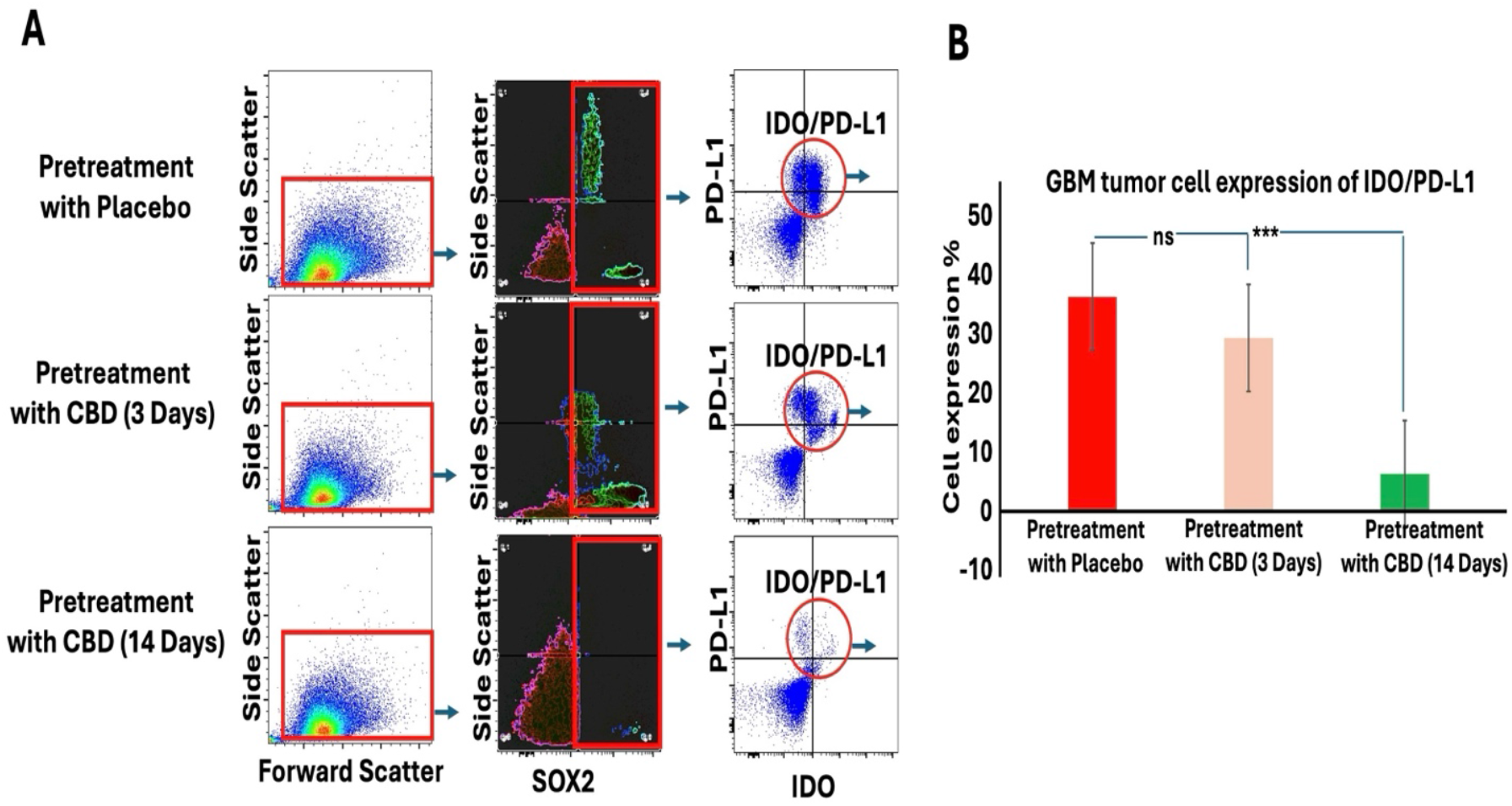
Inhaled CBD Pretreatment Decreases Immune Evasion Signals in GBM. **A)** Flow cytometry analysis was used to assess the expression of immune evasion signals, IDO and PD-L1, in GBM tumors following 14 days of inhaled CBD pretreatment. Tumor cells were identified based on forward light scatter (FSC) and side light scatter (SSC) (left panels), followed by gating for SOX2 expression, a marker of stem-like properties (middle panels). Dot plots show the reduction of IDO and PD-L1 expression specifically in SOX2-positive cells following 14-day CBD pretreatment. **B)** Quantification of IDO and PD-L1 expression in SOX2-positive cells revealed a significant decrease in both markers after 14-ay CBD pretreatment (p < 0.001), compared to the placebo and 3-day CBD groups. These results suggest that prolonged CBD pretreatment reduces immune evasion signals in GBM, particularly in stem-like tumor cells, potentially enhancing anti-tumor immunity.

### Inhaled CBD pretreatment reduced MGMT and Ki-67 expression in GBM tumors

Immunofluorescence staining and quantification revealed that mice pretreated with inhaled CBD for 14 days exhibited significantly reduced expression of MGMT and Ki-67 compared to both the 3-day CBD and placebo groups (p < 0.001) (Figure 4A-B). MGMT is a DNA repair enzyme associated with resistance to alkylating chemotherapy agents, such as Temozolomide (TMZ), and elevated levels are typically indicative of poor therapeutic response [19–20]. Ki-67, a marker of cellular proliferation, is strongly correlated with tumor cell growth, aggressiveness, and poor prognosis [21]. These findings suggest that CBD pretreatment may modulate key biomarkers involved in GBM progression, potentially enhancing the tumor’s sensitivity to treatment and improving therapeutic outcomes. To further confirm these results, we performed Western blotting analysis for MGMT. The Western blot data corroborated the immunofluorescence results, showing a consistent reduction in MGMT protein expression in the 14-day CBD pretreatment group compared to the placebo and 3-day CBD groups (Figure 4C). This additional analysis reinforces the notion that prolonged CBD exposure effectively modulates key biomarkers associated with GBM progression, supporting the potential for CBD to enhance therapeutic efficacy in GBM treatment.

**Figure 4.**
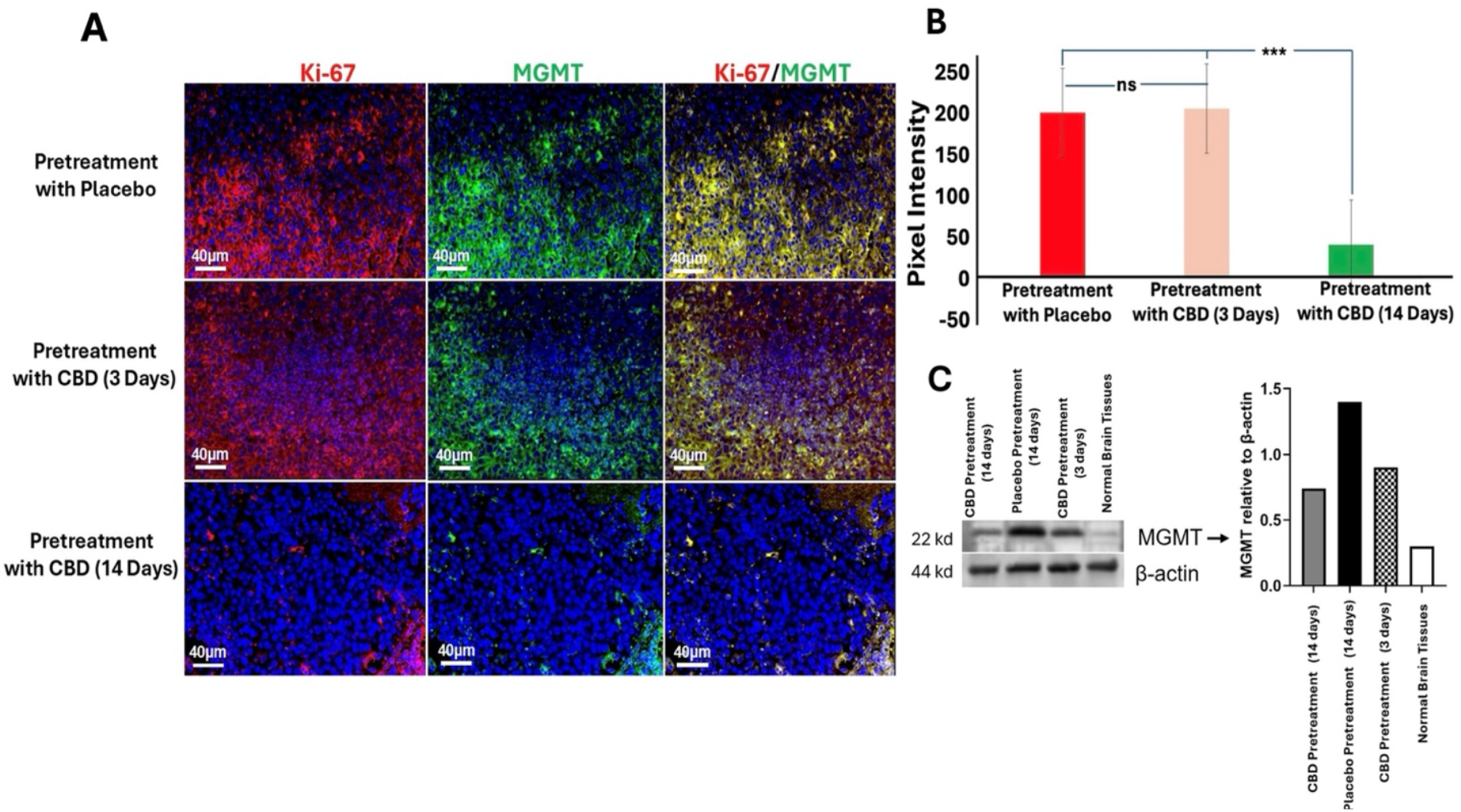
Inhaled CBD Pretreatment Reduces MGMT and Ki-67 in GBM Tumorsa) **A)** Immunofluorescence staining of MGMT and Ki-67 in GBM tumors from mice pretreated with inhaled CBD for 14 days, 3 days, or placebo. Representative images show that MGMT and Ki-67 levels were significantly reduced in the 14-day CBD pretreatment group compared to both the 3-day CBD and placebo groups (Scale bar: 40 μm). **B)** Quantification of staining intensity revealed significantly lower levels of MGMT and Ki-67 in the 14-day CBD pretreatment group compared to the 3-day CBD and placebo groups (p < 0.001). These results suggest that CBD pretreatment reduces key biomarkers associated with chemotherapy resistance and tumor proliferation, potentially enhancing therapeutic sensitivity. **C)** To further validate the effect of CBD on tumor progression, we assessed MGMT expression levels using Western blotting. The blot shows increased MGMT expression in the GBM tumors of mice pretreated with placebo (lane 2) compared to the normal brain tissue used as a negative control (lane 4). Long-term CBD pretreatment (14 days, lane 1) significantly reduced MGMT expression in the tumor compared to acute CBD pretreatment (3 days, lane 3), corroborating the reduction observed in the immunofluorescence analysis.

## Discussion

This study demonstrates that prolonged pretreatment with inhaled CBD (14 days) significantly inhibits glioblastoma (GBM) tumor growth in a murine model, whereas shorter pretreatment durations (3 days) or placebo treatments failed to show the same effect. Several key findings support this observation: (1) a notable reduction in tumor volume in the 14-day CBD pretreatment group compared to the placebo and 3-day CBD groups, (2) a decrease in the expression of SOX2, a marker of stem-like properties, along with immune checkpoint regulators IDO and PD-L1 in the 14-day CBD pretreatment group, and (3) a reduction in MGMT and Ki-67, biomarkers associated with DNA repair and cellular proliferation, respectively.

Importantly, the consistency between bioluminescent imaging and anatomical tumor quantification underscores the robustness of the anti-tumor effect observed with prolonged CBD pretreatment. While individual variability in photon flux patterns was noted—such as flat or modest BLI increases in some mice—this did not preclude a clear histological and volumetric confirmation of tumor suppression. These findings emphasize the need to interpret BLI data in the context of anatomical validation. Moreover, the improved survival seen in the prolonged CBD group further supports the biological relevance of these tumor growth differences. Although detailed imaging and molecular analyses were derived from a representative cohort, this approach ensured consistency in longitudinal data collection and was supported by survival trends replicated across independent experimental cohorts.

The most striking outcome of these findings is the inhibition of GBM growth following prolonged CBD pretreatment, which holds promise not only as a potential therapeutic strategy but also in slowing tumor progression, a critical challenge in GBM treatment [22]. CBD’s early intervention in modulating the tumor microenvironment and molecular pathways could help reduce the tumor cells’ ability to evade immune surveillance, suppress tumor proliferation, and overcome resistance mechanisms. This is particularly significant post-surgery, where CBD pretreatment could help mitigate tumor resistance to therapy, enhance tumor control, and reduce the risk of recurrence. Thus, CBD pretreatment has the potential to improve both survival rates and the overall quality of life for patients, particularly by reducing recurrence and improving chemotherapy efficacy through the regulation of resistance mechanisms like MGMT.

CBD’s favorable safety profile and lack of observed adverse effects—including weight loss, respiratory irritation, or behavioral changes during the 14-day aerosol exposure— further support its candidacy as a prophylactic agent. Preclinical studies have shown that CBD is well-tolerated, with minimal toxicity and no psychoactive effects, even at relatively high doses. Additionally, its ability to modulate oxidative stress, reduce neuroinflammation, and protect neural integrity aligns with its potential preventive role in neurological malignancies like GBM. These attributes offer a strong translational rationale for early administration in high-risk or pre-symptomatic settings, where safety and tolerability are paramount.

The inhalation route of CBD administration offers unique advantages, especially in the context of GBM treatment. While cannabinoids have primarily been investigated through oral or systemic routes, inhalation ensures rapid absorption, improved bioavailability, and enhanced blood-brain barrier (BBB) penetration, making it an ideal non-invasive approach for targeting brain tumors [23–25]. Although direct comparative data across routes are limited in GBM, prior studies suggest inhaled CBD outperforms oral forms in other neurological contexts [26]. This evidence supports our rationale for selecting the inhalation route, as briefly discussed earlier in the study. Furthermore, the duration-dependent effect observed in this study highlights the necessity of prolonged CBD exposure to achieve therapeutic effects. The results suggest that 14 days of pretreatment are crucial for modulating key molecular pathways involved in tumor growth and survival, marking a significant finding for optimizing treatment strategies.

The regulation of SOX2-positive cells is another critical outcome of this study. SOX2-positive cells are known for their stem-like properties and play a significant role in tumor resistance and recurrence in GBM [27]. By reducing the expression of immune checkpoint molecules such as IDO and PD-L1 in these cells, CBD pretreatment directly targets the most aggressive and therapy-resistant subpopulation of GBM cells. IDO and PD-L1 are essential for immune evasion and immune suppression, and downregulating these markers could enhance the immune system’s ability to recognize and attack tumor cells, thereby improving treatment efficacy [28–29]. Additionally, the regulation of MGMT and Ki67 further reinforces the potential of CBD as a therapeutic strategy. MGMT, known to confer resistance to TMZ, was significantly reduced following CBD pretreatment, suggesting that CBD could modify GBM tumors’ resistance to TMZ, improving its chemotherapeutic efficacy [19]. Similarly, Ki67, a key marker of cell proliferation, was downregulated by CBD, suggesting that it may reduce the tumor’s proliferative capacity [21]. Prior research has demonstrated that CBD can downregulate MGMT expression in glioma cells, particularly in vitro, through mechanisms involving oxidative stress and modulation of epigenetic pathways [30–31]. These findings provide a rationale for our current hypothesis that similar regulatory mechanisms may underlie the observed MGMT suppression in vivo following inhaled CBD pretreatment.

The CBD-induced reduction of these pivotal biomolecules including MGMT, IDO, PD-L1 and Ki67 may be well due to counter inflammatory and regulatory nature of CBD. CBD is reported to downregulate NF-κB and STAT3 signaling pathways which play central roles in the inflammatory process, immune checkpoint expression, and oncogenic transcription [32]. Additionally, CBD influences endoplasmic reticulum stress and oxidative stress responses in tumor cells, which may suppress tumor-promoting cytokine production and cell cycle progression. Altogether, these interactions may provide reasonable hypothetical rational to explanation the impact of CBD-pretreatment on the immune evasion, proliferation, and DNA repair markers in this mouse model of GBM. Together with the reduction in PD-L1 and IDO in SOX2-positive cells, these results highlight CBD’s multifaceted approach in targeting immune evasion, tumor growth, and treatment resistance, offering a comprehensive strategy to enhance the efficacy of existing therapies and improve long-term outcomes for GBM patients.

This study opens new avenues for research into pretreatment strategies for GBM, a largely underexplored area. The success of inhaled CBD in this model lays the foundation for further exploration of other cannabinoids or related compounds that may exhibit similar anti-tumor effects. Future directions should include validating behavioral tolerance, stress responses, and cognitive outcomes following prolonged CBD aerosol exposure to strengthen translational relevance. Additionally, evaluating the optimal dosing and duration of CBD pretreatment will be crucial in maximizing therapeutic efficacy and minimizing potential side effects. Combining CBD pretreatment with chemotherapy or immunotherapy could also be explored as a synergistic approach to treating glioblastoma.

### Limitations

While our findings are promising, several limitations should be acknowledged. This study was conducted in a murine model, and results may not fully translate to human patients, underscoring the need for clinical trials to evaluate the safety and efficacy of inhaled CBD in glioblastoma. The molecular mechanisms underlying CBD’s anti-tumor effects remain incompletely understood, and future studies should further investigate its interactions with cannabinoid receptors, oxidative stress pathways, and epigenetic modulators. Optimal dosing and duration of CBD pretreatment also require careful refinement to maximize therapeutic benefit while minimizing side effects. Although we briefly addressed pharmacokinetics and bioavailability of inhaled CBD in the Methods section, the lack of direct comparisons across delivery routes within this GBM model limits interpretation; ongoing work in our lab aims to address this. While bioluminescent imaging and related outcomes were derived from a representative cohort selected for complete survival to permit consistent imaging, survival analysis was performed across two independent cohorts to improve statistical power. Finally, although our group sizes (n=5 per group, replicated across two cohorts) reflected consistent trends, larger-scale studies will be essential to further validate and generalize these findings.

## Conclusion

This preclinical investigation demonstrates that prolonged inhaled CBD pretreatment significantly suppresses glioblastoma (GBM) progression in a murine model by targeting multiple hallmarks of tumor biology. Key outcomes include reduced tumor burden and modulation of molecular markers associated with tumor stemness (SOX2), immune evasion (PD-L1, IDO), cellular proliferation (Ki67), and chemoresistance (MGMT). These effects were accompanied by immunomodulatory and antiproliferative shifts in the tumor microenvironment, consistent with a transition toward a less aggressive phenotype. Notably, the superiority of a 14-day pretreatment regimen over shorter durations underscores the importance of sustained exposure and supports inhalation as a promising delivery route for central nervous system therapeutics. Collectively, these findings support the development of CBD as a non-invasive, prophylactic adjunct to standard GBM treatments and provide a strong rationale for further translational studies aimed at optimizing CBD-based interventions to improve clinical outcomes in this aggressive malignancy.

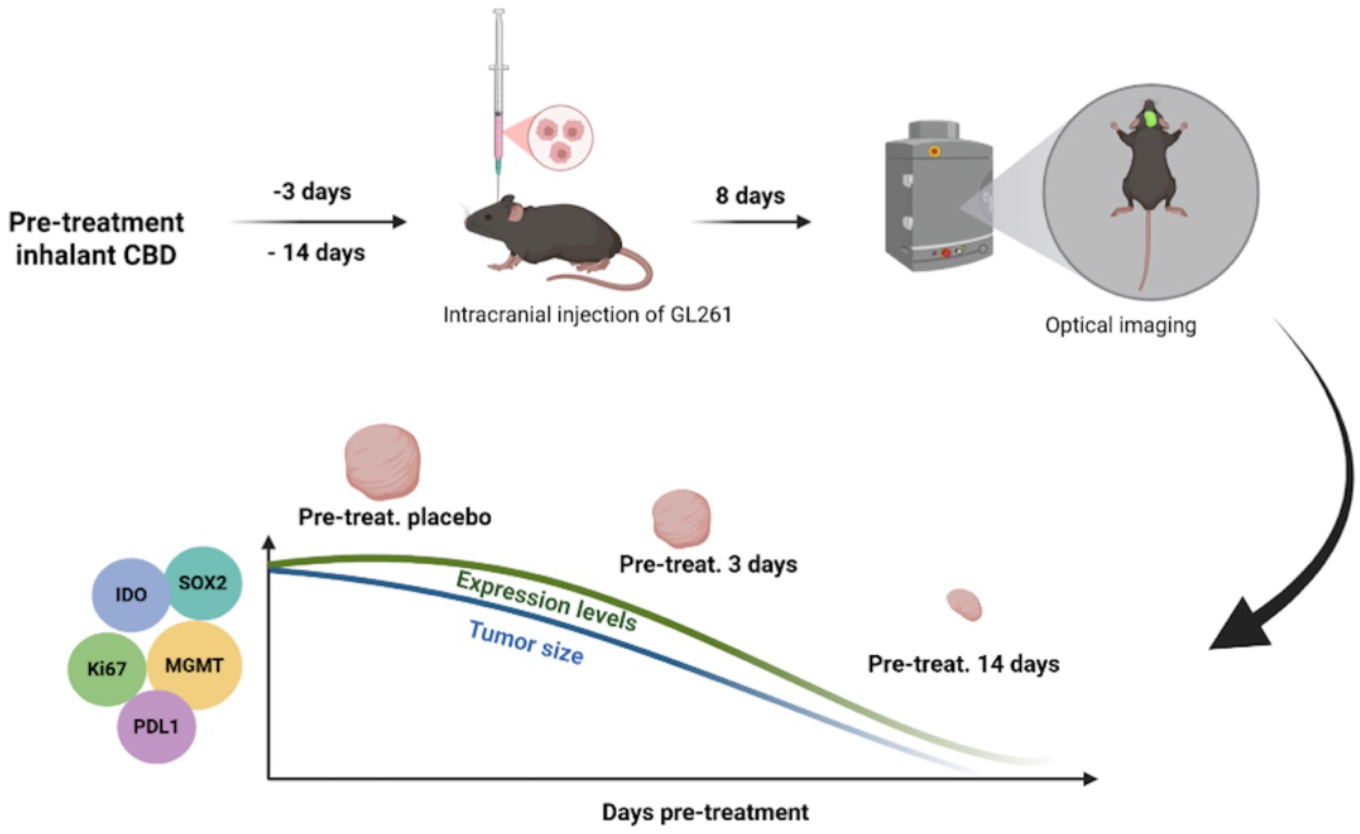

## Statements & Declarations

## Acknowledgements

Authors are thankful to Thriftmaster Holding Group for providing the inhalant CBD for this study. Authors also thank Medicinal Cannabis of Georgia for providing help in the analysis, processing the data and optimizing the CBD dosage.

## Declaration of AI Assistance

AI-based software (e.g., ChatGPT) was used solely for language refinement and grammar editing; all scientific content and interpretation were entirely developed by the authors.

## Author contributions

LPW and BB: Study conception and design, analysis and interpretation of data, and drafting the article., NPM, ELS, SEN, BIB, HMR, AK, and BH: Acquisition of data and drafting the article., HMR, NY, GW, FLV, JCY, SA, and ASA: Editing and Scientific Contribution., LPW, BB and ASA: Resourcing and management.

## Funding

This work was supported by institutional seed funding from the Dental College of Georgia at Augusta University.

## Competing interests

(1) Lei Phillip Wang, Babak Baban, and Jack Yu are members of Medicinal Cannabis of Georgia with no financial interest. (2) All other authors declare no conflict of interest. (3)Thriftmaster Holding Group (THG) is the provider of CBD inhalers and has a licensing contract with Augusta University. (4) THG had no role in study design, data collection and analysis, decision to publish, or preparation of the manuscript.

## Data availability

Data is provided within the manuscript or supplementary information files.

## Ethics approval

Animal experiments were approved by the Institutional Animal Care and Use Committee (IACUC, Protocol#2011-0062) of Augusta University and followed the IACUC guidelines. There was no human subjects/samples used in this study.

## References

1- Wang LP, Chagas PS, Salles ÉL, Naeini SE, Gouron J, Rogers HM, Khodadadi H, Bhandari B, Alptekin A, Qin X, Vaibhav K, Costigliola V, Hess DC, Dhandapani KM, Arbab AS, Rutkowski MJ, Yu JC, Baban B. Altering biomolecular condensates as a potential mechanism that mediates cannabidiol effect on glioblastoma. Med Oncol. 2024 May 7;41(6):140.

2- Ostrom QT, Price M, Neff C, Cioffi G, Waite KA, Kruchko C, Barnholtz-Sloan JS. CBTRUS Statistical Report: Primary Brain and Other Central Nervous System Tumors Diagnosed in the United States in 2015-2019. Neuro Oncol. 2022 Oct 5;24(Suppl 5):v1–v95.

3- Khodadadi H, Salles ÉL, Alptekin A, Mehrabian D, Rutkowski M, Arbab AS, Yeudall WA, Yu JC, Morgan JC, Hess DC, Vaibhav K, Dhandapani KM, Baban B. Inhalant Cannabidiol Inhibits Glioblastoma Progression Through Regulation of Tumor Microenvironment. Cannabis Cannabinoid Res. 2023 Oct;8(5):824–834.

4- Becker AP, Sells BE, Haque SJ, Chakravarti A. Tumor Heterogeneity in Glioblastomas: From Light Microscopy to Molecular Pathology. Cancers (Basel). 2021 Feb 12;13(4):761.

5- Fisher JP, Adamson DC. Current FDA-Approved Therapies for High-Grade Malignant Gliomas. Biomedicines. 2021 Mar 22;9(3):324.

6- Angom RS, Nakka NMR, Bhattacharya S. Advances in Glioblastoma Therapy: An Update on Current Approaches. Brain Sci. 2023 Oct 31;13(11):1536.

7- Osuka S, Van Meir EG. Overcoming therapeutic resistance in glioblastoma: the way forward. J Clin Invest. 2017 Feb 1;127(2):415–426.

8- Waqar M, Trifiletti DM, McBain C, O’Connor J, Coope DJ, Akkari L, Quinones-Hinojosa A, Borst GR. Early Therapeutic Interventions for Newly Diagnosed Glioblastoma: Rationale and Review of the Literature. Curr Oncol Rep. 2022 Mar;24(3):311–324.

9- Müller Bark J, Kulasinghe A, Chua B, Day BW, Punyadeera C. Circulating biomarkers in patients with glioblastoma. Br J Cancer. 2020 Feb;122(3):295–305.

10- Dumitru CA, Sandalcioglu IE, Karsak M. Cannabinoids in Glioblastoma Therapy: New Applications for Old Drugs. Front Mol Neurosci. 2018 May 16;11:159.

11- Volmar MNM, Cheng J, Alenezi H, Richter S, Haug A, Hassan Z, Goldberg M, Li Y, Hou M, Herold-Mende C, Maire CL, Lamszus K, Flüh C, Held-Feindt J, Gargiulo G, Topping GJ, Schilling F, Saur D, Schneider G, Synowitz M, Schick JA, Kälin RE, Glass R. Cannabidiol converts NF-κB into a tumor suppressor in glioblastoma with defined antioxidative properties. Neuro Oncol. 2021 Nov 2;23(11):1898–1910.

12- Seltzer ES, Watters AK, MacKenzie D Jr, Granat LM, Zhang D. Cannabidiol (CBD) as a Promising Anti-Cancer Drug. Cancers (Basel). 2020 Oct 30;12(11):3203.

13- Cleirec G, Desmier E, Lacatus C, Lesgourgues S, Braun A, Peloso C, Obadia C. Efficiency of Inhaled Cannabidiol in Cannabis Use Disorder: The Pilot Study Cannavap. Front Psychiatry. 2022 May 24;13:899221.

14- Soroceanu L, Singer E, Dighe P, Sidorov M, Limbad C, Rodriquez-Brotons A, Rix P, Woo RWL, Dickinson L, Desprez PY, McAllister SD. Cannabidiol inhibits RAD51 and sensitizes glioblastoma to temozolomide in multiple orthotopic tumor models. Neurooncol Adv. 2022 Feb 17;4(1):vdac019.

15- Brookes A, Kindon N, Scurr DJ, Alexander MR, Gershkovich P, Bradshaw TD. Cannabidiol and fluorinated derivative anti-cancer properties against glioblastoma multiforme cell lines, and synergy with imidazotetrazine agents. BJC Rep. 2024 Sep 9;2(1):67.

16- Lucas CJ, Galettis P, Schneider J. The pharmacokinetics and the pharmacodynamics of cannabinoids. Br J Clin Pharmacol. 2018 Nov;84(11):2477–2482.

17- Devinsky O, Kraft K, Rusch L, Fein M, Leone-Bay A. Improved Bioavailability with Dry Powder Cannabidiol Inhalation: A Phase 1 Clinical Study. J Pharm Sci. 2021 Dec;110(12):3946–3952.

18- Al-Shabrawey M, Mussell R, Kahook K, Tawfik A, Eladl M, Sarthy V, Nussbaum J, El-Marakby A, Park SY, Gurel Z, Sheibani N, Maddipati KR. Increased expression and activity of 12-lipoxygenase in oxygen-induced ischemic retinopathy and proliferative diabetic retinopathy: implications in retinal neovascularization. Diabetes. 2011 Feb;60(2):614–24.

19- Kitange GJ, Carlson BL, Schroeder MA, Grogan PT, Lamont JD, Decker PA, Wu W, James CD, Sarkaria JN. Induction of MGMT expression is associated with temozolomide resistance in glioblastoma xenografts. Neuro Oncol. 2009 Jun;11(3):281–91.

20- Yu W, Zhang L, Wei Q, Shao A. O6-Methylguanine-DNA Methyltransferase (MGMT): Challenges and New Opportunities in Glioma Chemotherapy. Front Oncol. 2020 Jan 17;9:1547.

21- Li LT, Jiang G, Chen Q, Zheng JN. Ki67 is a promising molecular target in the diagnosis of cancer (review). Mol Med Rep. 2015 Mar;11(3):1566–72.

22- Vaz-Salgado MA, Villamayor M, Albarrán V, Alía V, Sotoca P, Chamorro J, Rosero D, Barrill AM, Martín M, Fernandez E, Gutierrez JA, Rojas-Medina LM, Ley L. Recurrent Glioblastoma: A Review of the Treatment Options. Cancers (Basel). 2023 Aug 26;15(17):4279.

23- Kolesarova M, Simko P, Urbanska N, Kiskova T. Exploring the Potential of Cannabinoid Nanodelivery Systems for CNS Disorders. Pharmaceutics. 2023 Jan 6;15(1):204.

24- Feng S, Pan Y, Lu P, Li N, Zhu W, Hao Z. From bench to bedside: the application of cannabidiol in glioma. J Transl Med. 2024 Jul 11;22(1):648.

25- Calapai F, Cardia L, Sorbara EE, Navarra M, Gangemi S, Calapai G, Mannucci C. Cannabinoids, Blood-Brain Barrier, and Brain Disposition. Pharmaceutics. 2020 Mar 15;12(3):265.

26- Bhandari B, Naeini SE, Rezaee S, Rogers HM, Khodadadi H, Bosomtwi A, Seyyedi M, MacKinnon NJ, Dhandapani KM, Salles ÉL, Hess DC, Yu JC, Moore-Hill D, Vale FL, Wang LP, Baban B. Optimization of seizure prevention by cannabidiol (CBD). Transl Neurosci. 2025 Mar 28;16(1):20220362.

27- Lopez-Bertoni H, Johnson A, Rui Y, Lal B, Sall S, Malloy M, Coulter JB, Lugo-Fagundo M, Shudir S, Khela H, Caputo C, Green JJ, Laterra J. Sox2 induces glioblastoma cell stemness and tumor propagation by repressing TET2 and deregulating 5hmC and 5mC DNA modifications. Signal Transduct Target Ther. 2022 Feb 9;7(1):37.

28- Parvez A, Choudhary F, Mudgal P, Khan R, Qureshi KA, Farooqi H, Aspatwar A. PD-1 and PD-L1: architects of immune symphony and immunotherapy breakthroughs in cancer treatment. Front Immunol. 2023 Dec 1;14:1296341.

29- Jiang X, Wang J, Deng X, Xiong F, Ge J, Xiang B, Wu X, Ma J, Zhou M, Li X, Li Y, Li G, Xiong W, Guo C, Zeng Z. Role of the tumor microenvironment in PD-L1/PD-1-mediated tumor immune escape. Mol Cancer. 2019 Jan 15;18(1):10.

30- Ladin DA, Soliman E, Griffin L, Van Dross R. Preclinical and Clinical Assessment of Cannabinoids as Anti-Cancer Agents. Front Pharmacol. 2016 Oct 7;7:361.

31- Brookes A, Kindon N, Scurr DJ, Alexander MR, Gershkovich P, Bradshaw TD. Cannabidiol and fluorinated derivative anti-cancer properties against glioblastoma multiforme cell lines, and synergy with imidazotetrazine agents. BJC Rep. 2024 Sep 9;2(1):67.

32- Peyravian N, Deo S, Daunert S, Jimenez JJ. Cannabidiol as a Novel Therapeutic for Immune Modulation. Immunotargets Ther. 2020 Aug 18;9:131–140.

